# Corticotropin-Releasing Factor Induces Functional And Structural Synaptic Remodelling In Acute Stress

**DOI:** 10.1101/2021.05.19.443320

**Authors:** Dorien Vandael, Keimpe Wierda, Katlijn Vints, Pieter Baatsen, Lies De Groef, Lieve Moons, Vasily Rybakin, Natalia V. Gounko

## Abstract

Biological responses to internal and external stress factors involve highly conserved mechanisms, using a tightly coordinated interplay of many factors. Corticotropin-releasing factor (CRF) plays a central role in organizing these lifesaving physiological responses to stress. We show that CRF rapidly and reversibly changes Schaffer Collateral input into hippocampal CA1 pyramidal cells (PC), by modulating both functional and structural aspects of these synapses. Host exposure to acute stress, *in vivo* CRF injection, and *ex vivo* CRF application all result in fast *de novo* formation and remodeling of existing dendritic spines. Functionally, CRF leads to a rapid increase in synaptic strength of Schaffer collateral input into CA1 neurons, e.g. increase in spontaneous neurotransmitter release, paired-pulse facilitation and repetitive excitability and improves long-term synaptic plasticity: LTP and LTD. In line with the changes in synaptic function, CRF increases the number of presynaptic vesicles, induces redistribution of vesicles towards the active zone increases active zone size, and improves the alignment of the pre- and post-synaptic compartments. Together, CRF rapidly enhances synaptic communication in the hippocampus, potentially playing a crucial role in the enhanced memory consolidation in acute stress.

## Introduction

Stress is a fundamental homeostatic reaction to any stimulus (1,2), which can biologically manifest itself as predominantly positive ‘eustress’ or predominantly as negative ‘distress’ (3). Acute stress is an instantaneous and precise reaction to internal and environmental factors (4–6). Although mechanisms involved in regulating stress responses are well documented for the hypothalamic-pituitary-adrenal (HPA) axis pathway, the effect of stress on other regions of the brain is still not well understood (7,8). Among the many hormones, neuropeptides, and mediators involved in the stress response, CRF stands out due to its dual systemic (hormonal) and central (neuromodulatory) roles (8–10). Centrally, CRF acts as a neuromodulator of synaptic transmission which can be rapidly and locally released and acts within milliseconds (7) by binding to two different G protein-coupled receptors: CRF-receptor (CRF-R) 1 and 2 (4,7,10). Activation of these receptors can result in a comprehensive array of cellular effects depending on the brain region and the specific CRF-family ligand binding (8,11). This can explain the diversity of responses reported in different brain regions to the same stressor. In the hippocampus, a region known for its involvement in learning and memory processes, CRF is expressed by GABAergic interneurons, which innervate PCs in CA1 and CA3 (12,13) and these PCs express CRF-Rs in distinct subcellular regions (4,14,15). The effects of stress on - hippocampus dependent - memory storage and consolidation are complex (4,16–18). Mild or short stress enhances hippocampal functioning by promoting synaptic strengthening and by augmenting frequency of mEPSCs and glutamate release probability (7,19), while profound and chronic stress has detrimental effects, manifesting in the reduction in dendritic complexity and spine density in the hippocampus (12). This spine loss is associated with attenuation of both long-term potentiation (LTP) and long-term depression (LTD), and correlates with reported memory defects (7,20–22). CRF contributes to the initiation of those stress induced neuronal changes (7,12,23,24) in a dose-, time- and context-dependent manner (4,16,25,26). Especially the period of CRF exposure can have crucial deferential effects on learning and memory processes and might result in opposite effects (4,16,25). For example, short-term CRF application increases LTP (27) while prolonged exposure impairs hippocampal LTP (28).

Previous studies on structural changes reported a decrease in spine number and reduction of dendritic complexity of PCs in CA1 and CA3 after long-term exposure to CRF (24,29,30). In addition, the underlying molecular pathways of CRF-dependent plasticity have been mostly studied in the presence of high CRF concentrations and using *in vitro* assays. However, the acute effect of CRF in a physiologically relevant concentration (<250 nm) (8,31,32) on synaptic architecture and function in the hippocampus remain elusive.

Here, we show that acute stress, CRF stereotactic injections *in vivo*, and application of CRF *ex vivo* induces spine maturation and increases spine density. At the synapse level, we demonstrate that acute CRF increases the presynaptic vesicular pool size, increases synapse number, induces a redistribution of synaptic vesicles towards the active zone and increases alignment of pre- and postsynaptic compartments. In line with these structural changes, we found that CRF facilitates synaptic transmission and increases synaptic reliability. In addition, CRF enhances long term synaptic plasticity, which requires reciprocal activation of both CRF-R receptors. Taken together, this study provides evidence that CRF is a crucial player in shaping the cellular response of hippocampal CA1 PCs during acute stress.

## Materials and Methods

### Animals

All animal experiments were approved by the KU Leuven Ethical Animal Welfare Committee (protocol P019/2017) and were performed following the Animal Welfare Committee guidelines of the KU Leuven, Belgium. Mice were housed in a pathogen-free facility under standard housing conditions. In total, 113 male C57BL/6Jax mice (P18-20), 24 male Thy1-YFP-H line, B6.Cg-Tg(Thy1-YFP)HJrs/J (P21-23, JAX 003782) and 4 male C57BL/6J-Tg(Thy1-GCaMP6)GP4.12Dkim/J (P18-20, JAX 028278) were used.

### Acute stress induction and stereotactic injections in vivo

Thy1-YFP-H mice were used for acute stress and stereotactic injections with 100nM CRF. For acute stress, we used two paradigms: foot shock (FS) and predator odor (PO) (33,34). For PO, mice were transferred from their home cage to a clean cage and subsequently exposed to either PO (domestic cat urine/fur mixed with cotton wool) or ambient air (cotton wool, control) (35). The FS was performed as described before (36). Briefly, control animals stayed in the home cage without any handling. Acute stress FS protocol was a 0.1mA electrical stimulation for 2 seconds. 20 minutes after the stimulus, mice were deeply anesthetized with a mixture of ketamine/xylazine and cardiac puncture was carried out for trunk blood collection. Blood plasma was stored for corticosterone (CORT) ELISA analysis. Brains were collected after trancardiac perfusion with 4% paraformaldehyde (PFA; EMS) in 0.1M phosphate buffer (PB; EMS). From each animal, one hemisphere was used for spine analysis of the PCs dendrites in the proximal region of CA1-Stratum Radiatum (SR), the other hemisphere was used for *cfos* and corticotropin-releasing hormone *(crh)* mRNA *in situ* hybridization (ISH) experiments. All acute stress experiments and blood collection were done during the same time of day (controlled for circadian rhythm).

For stereotactic injections of CRF in PCs CA1 hippocampus, mice were anesthetized by isoflurane and placed in a stereotactic frame with sustained anesthesia during and post injection. 300nl of 100nM CRF with a rate of 10nl/sec was unilateral injected using a Nanoject II Auto-Nanoliter Injector (Drummond) using stereotactic coordinates: AP-2, ML-1.8, D-1.5 mm. The other (non-injected) hemisphere was used as a control (37). Animals were perfused with 4% PFA in 0.1M PB, 20 minutes after the injection. Until sample collection, animals were kept constantly under anesthesia. Brains were post-fixed at 4°C overnight. The following day, 100μm-thick vibratome sections were made and used for further processing (see below).

### Determination of hormone concentrations and ISH after acute stress

Plasma was separated from whole blood and stored at −80°C until further sample processing. CORT plasma levels were quantified using a CORT ELISA kit (DE4164, Demeditec Diagnostics). Blood plasma was 1:20 diluted with standard 0 solutions. Absorbance was determined at 450nm (reading) and 620-630nm (background subtraction) with a microtiter plate reader.

Basescope hybridization was performed with the Basescope Detection Reagent Kit v2-RED (Advanced Cell Diagnostics). Briefly, 14μm-thick cryosections of fixed frozen Thy1-YFP-H hemispheres of control and stressed mice were made. Superfrost slides (Thermofisher) with sections were baked at 60°C for one hour before dehydrating steps of ethanol. After pretreatment solution steps, sections were incubated with custom-synthesized Basescope probes (*cfos*, BA-Mm-Fos-3zz-st targeting 676-801 of NM_010234.3 or *crh*, BA-Mm-Crh-3zz-st targeting 752-893 of NM_205769.3) each targeting all predicted transcript variants, followed by amplifying hybridization processes. Between amplification steps, slices were washed with wash buffer. Finally, slides were incubated with Fast Red for 10 minutes at room temperature in the dark and counterstained with 50% hematoxylin before drying at 60°C. Brightfield images were taken with a Marzhauser Express 2 slide scanner (Nikon) using a 20X objective. After imaging, the layer of PCs CA1 from each section was used for probe quantification. Probe-positive areas and physical CA1 PC areas were manually segmented using Microscope Image Browser (MIB) (University of Helsinki) (38). Data have been expressed as probe-positive areas relative to PCs-occupied areas.

### Dendritic spine filling ex vivo

For dye filling experiments in hippocampal acute slices, C57BL/6Jax mice were used, as described (39). Briefly, animals were anaesthetized using isoflurane. After decapitation, the brain was quickly removed and transferred into ice-cold cutting solution: 83µM NaCl, 2.5mM KCl, 1mM NaH_2_PO_4_, 22mM glucose, 26.2mM NaHCO_3_, 0.5mM CaCl_2_, 3.3mM MgSO_4_, 72mM sucrose (Sigma), pH7.4 with 5% CO_2_/95% O2. 300μm coronal slices were cut with a Leica VT1200 vibratome. Slices could recover in a 34°C cutting solution for 35 minutes and for 30 minutes at room temperature (RT) prior to transfer into artificial cerebrospinal fluid (aCSF): 119mM NaCl, 2.5mM KCl, 1mM NaH_2_PO_4_, 26mM NaHCO_3_, 4mM MgCl_2_, 4mM CaCl_2_, 11mM glucose at pH7.4 with 5% CO_2_/ 95% O_2_. Glass borosilicate recording pipettes (resistance 3.5-5.5MΩ) were filled with 10mM Alexa 568 (Life Technologies) dissolved in internal solution: 15mM CsMSF, 20mM CsCl, 10mM HEPES, 2.5mM MgCl_2_, 4mM ATP, 0.4mM GTP, 10mM creatine phosphate and 0.6mM EGTA (Sigma Aldrich). Whole-cell configuration was used to fill CA1 PCs for 10-15 minutes in control slices and slices incubated with 100nM CRF added to the aCSF for 20 minutes. Hence, slices are incubated 10 minutes prior to the filling with aCSF and CRF. Treatment with blockers was carried out by directly adding them to the aCSF minimal 20 minutes before reaching whole cell mode. For condition of blockers with CRF, CRF was added 10 minutes after slices were exposed to the specific CRF-R blockers. Sections were fixed with 4% PFA and 2% sucrose in 0.1M PB at 4°C overnight.

### Spine imaging and analysis ex vivo and in vivo

After 4% PFA fixation overnight, brain slices were washed three times with 0.1 M PB and mounted using mounting medium (Vectashield). 100μm-thick vibratome sections were made from brains collected after acute stress paradigms and stereotactic injections of CRF, as described above. Secondary and tertiary dendrites of PCs in the proximal region of the CA1 were imaged with a Structured Illumination Microscopy (Elyra S.1, Zeiss) with a 63X plan-apochromat 1.4 oil DIC objective. Images were processed using the Zeiss software. Dendritic protrusions were counted in Z-stack (Z-step of 0.025µm) and quantified using ImageJ (NIH). We classified 5 spine types. Mushroom spines: possess a spine head of more than 0.5µm. Stubby: length shorter than 1.0 µm. Spine head diameter larger than spine length. Thin: length shorter than 1.0 µm possessing, spine head diameter shorter than spine length. Long thin: length between 1.0 and 1.5 µm. Filopodia: longer than 1.5µm.

### Electrophysiological and multi electrode array (MEA) ex vivo studies

#### Ex vivo

Acute slices (300μm) were prepared from C57BL/6Jax mice the same way as for *ex vivo* spine fillings, as described before (39). After recovery, brain slices were continuously perfused in a submerged chamber (Warner Instruments) at a rate of 3-4 ml/minutes with aCSF at pH7.4 with 5% CO_2_/ 95% O_2_. Control slices and slices incubated with 100nM CRF added to the aCSF for ∼20 minutes before recording were used. For mEPSCs, coronal sections were prepared and 1µM tetrodotoxin (TTX) was added to the aCSF. For paired-pulse recordings, train stimulation, and AMPA/NMDA characterization, sagittal slices were used and 20μM bicuculline was added to the aCSF. Whole-cell patch-clamp recordings were done using borosilicate glass recording pipettes (resistance 3.5-5.5MΩ) filled with a CsMSF-based internal solution (see *ex vivo* spine filling). Spontaneous input to CA1 PCs was recorded by whole-cell voltage-clamp recordings (Vm=−70mV and Rs compensation was set at ∼70%) from visually identifiable CA1 PCs, using a Multiclamp 700B amplifier (Axon Instruments) and analyzed using Mini Analysis program (Synaptosoft). For evoked recordings (Vm=-70 mV, Rs compensation ∼70%), Schaffer collaterals were stimulated using A-M systems 2100 isolation pulse stimulator and a 2-contact cluster microelectrode (CE2C55, FHC) placed in SR at the border of CA1-CA2. For paired-pulse ratio analysis, paired extracellular stimulations (interstimulus interval (ISI): 25, 50, 100, 200, 400, and 1000ms) were delivered every 20 seconds (each ISI was repeated 3 times) and peak amplitudes were calculated as the EPSC2/EPSC1 ratio. For train stimulations, 200 stimuli were delivered at the following frequencies: 2Hz, 5Hz, 10Hz, and 20Hz. Peak amplitudes and total charge were quantified and normalized to the first evoked response of the train. Peak AMPAR-mediated evoked EPSCs were measured in whole-cell voltage-clamp at a holding potential of −60mV, while the NMDAR-mediated component was measured 100ms after initiation of the combined AMPAR-and NMDAR-mediated EPSCs recorded at +40mV. Measurements were performed in a minimum of three independent preparations.

#### MEA

Parasagittal slices (300 μm) were prepared from C57BL/6Jax mice and used for fEPSPs recording using commercially available MEAs, 60 electrodes in an 8×8 lay-out (MEA2100, Multi Channel Systems) as described before (40,41). The recording chamber was perfused with aCSF and maintained at 32°C. A slice grid was put on the top of the slices to assure immobilization and optimal contact with electrodes. Data streams were sampled at 10 kHz. For each slice, a single electrode located underneath the Schaffer collateral pathway was visually selected for stimulation. Biphasic, constant voltage pulses (100µs pulse width) were applied to evoke fEPSPs from the Schaffer collaterals (SC) in the CA1. After establishing stable fEPSP signals (after approximately 30 minutes), an input/output curve was generated using stimulation intensities from 0.5 to 2.750V (in steps of 0.25V), each applied twice with 30-120 seconds interval was established. The stimulus intensity eliciting 35% of the maximal fEPSP amplitude was used for further stimulation.

Next, we recorded baseline fEPSPs for approximately 25 minutes (3 stimulations 15 seconds apart, every 3 minutes). For CRF conditions, after 5 minutes of baseline, we switched to aCSF with 100nM CRF, recorded 15 minutes of baseline, switched back to aCSF which normalized a stable baseline comparable to before CRF application. After reestablishing a stable baseline, we either applied train stimulations (LTP) or low frequency stimulations (LTD). LTP was introduced by three trains of high-frequency stimulation at 100Hz (100 stimuli at 100Hz), with 5 minutes interval. For induction of LTD, low frequency stimulation of 1Hz, 900 pulses was induced to introduce LTD in the CA1 region. Post-LTD or -LTP induction, fEPSPs were recorded for 65 minutes (3 stimulations 15 seconds apart, every 3 minutes).

### Calcium imaging in vivo

Acute coronal slices (300μm) were prepared from Thy1-GCaMP6 mice (see above). After recovery, brain slices were continuously perfused with aCSF during the imaging of the CA1 at RT with a two-photon system (VIVO 2-Photon platform, Intelligent Imaging Innovations GmbH) using a 20X objective. Imaging started in aCSF capturing 300 images of the region of interest (ROI), average of 15 frames per image, 30ms intervals. 600 images were taken: 300 control aCSF images and another 300 images where CRF was present in the aCSF. After 15 minutes with CRF in aCSF, another 600 images were taken with the same settings in the same ROI.

### Electron microscopy (EM) and analysis

Acute coronal slices (300µm) were prepared from C57BL/6Jax mice (see above). After recovery, control and CRF-treated slices (100nM CRF for 20 minutes) were fixed for at least 2 hours at room temperature. For synaptic morphology we used 4% PFA, 2% glutaraldehyde (EMS, USA), 0.2% picric acid (EMS, USA) in 0.1M PB, pH7.4 For active zone (AZ) and postsynaptic densities (PSD) quantification, we used 4% PFA in 0.1M PB, pH7.4.

For synaptic morphology analysis with transmission electron microscopy (TEM), after fixation slices were subsequently washed with 0.1M PB and 0.1M cacodylate buffer and post-fixed for 60 minutes on ice in 0.1M cacodylate buffer (EMS, USA) containing 1% OsO_4_ (EMS, USA) and 1.5% C_6_FeK_4_N_6_ (EMS, USA), pH 7.6. Next, slices were washed once with 0.1M cacodylate buffer, and then with dH_2_O. The slices were contrasted with 0.5% uranyl acetate (EMS, USA) in 25% methanol at 4°C overnight. The following day, slices were washed with dH_2_O and stained on bloc with Walton’s lead aspartate (39) at 60°C for 30 minutes, and washed with dH_2_O. Afterwards, the samples were dehydrated in a graded series of ethanol solutions and were treated twice for 10 minutes with propylene oxide and infiltrated with medium Epon 812/propylene oxide mixtures. The next day, sections were flat embedded in medium composition of Epon 812 (EMS, USA) between two microscopic slides and ACLAR film (EMS) and polymerized for 2 days at 60°C.

For visualization and analysis of AZ and PSD with TEM and focused ion beam scanning electron microscope (FIB-SEM), After fixation slices were washed with 0.1M PB. and dehydrated in a graded series of ethanol solutions. Afterward, slices were treated for 30 minutes at 60°C in 1% ethanolic phosphotungstic acid (PTA; MP Biomedicals). Slices were washed with pure ethanol and subsequently with pure acetone. The slices were contrasted with 2% uranyl acetate in acetone at 60°C for 20 minutes. Slices were then washed with acetone and incubated in 0.5% lead acetate in acetone at 60°C for 20 minutes, washed with acetone and infiltrated with hard Epon 812/acetone mixtures. The next day, slices were embedded in hard composition of Epon 812 and polymerized for 2 days at 60°C.

For TEM imaging ultrathin sections (70 nm) were collected on single slot copper grids and counterstained with uranyl acetate and lead citrate. Images of these sections were made at 25kX magnification for synaptic boutons morphology and at 15kX magnification for AZ/PSD analysis, using a TEM (JEM1400, Jeol) equipped with a SIS Quemesa (Olympus) camera operated at 80kV.

For the FIB-SEM, the embedded samples were coated with ∼8 nm platinum. FIB-SEM imaging is performed using a Crossbeam 540 (Zeiss) system with Atlas 3D software. The FIB-SEM was used to remove a 5nm-thick layer by propelling gallium ions at the surface of the specimen. Image acquisition was done at 1.5kV (0.005 µm/pixel) using a backscattered electron detector, at 5kX magnification. Images were aligned with Atlas 3D software.

PCs CA1 synapses were identified by their morphology and localization. Image segmentation of individual pre- and postsynaptic terminals, PSDs, AZs and synaptic vesicles was performed initially by using MIB software. For vesicle analysis, we estimated the shortest straight path connecting the center of vesicle to the AZ and calculated the smallest angles between the directions of this path. The statistics for synaptic surface area, AZ/PSD area and length, number of vesicles and distance from AZ was collected using a custom-made script in ImageJ. Amira software was used for visualization of AZs and PSDs in 3D.

### Drugs and treatments

The used dilutions: Alexa 568 hydrazide - 10 mM (Thermo Fisher Scientific), Antisauvagine-30 (aSvg) - 150 nM (Tocris), bicuculline - 20 μM (Sigma Aldrich), CRF - 100 nM (Bachem), NBI 27914 (NBI) - 1.2 µM (Tocris), TTX - 1 µM (Tocris). Besides Alexa568, all drugs were dissolved in DMSO prior added into used solutions.

### Quantification and statistical analysis

Data analysis was carried out in ImageJ (NIH), Clampfit (Molecular devices), MiniAnalysis (Synaptosoft), Multichannel analyzer software (Multi channel systems), Microscope Image Browser (MIB, University of Helsinki), Amira (Thermo Scientific), Atlas 3D (Zeiss) and Excel (Microsoft). Data statistic was carried out in GraphPad Prism 8 (GraphPad software).

We first evaluated the quantitative sample distributions for normality using the D’Agostino-Pearson test. Subsequently, either Mann-Whitney test (for non-normal distributions) or unpaired t-test (for normal distributions) was used to compare statistical differences between any two groups. Comparisons between multiple groups were performed with the Kruskal-Wallis analysis of variance (ANOVA) followed by Dunn’s multiple comparison test (for non-normal distributions) or with one-way ANOVA followed by Dunnett’s multiple comparison test (for normal distributions). Results were evaluated at a 5% significance level.

## Results

### Both short-term stress and CRF treatment induce spine formation in vivo

Previous studies in different brain regions have shown stress induces changes in spine density and morphology (42–45). To investigate the effect of short-term stress on spines of hippocampal CA1 PCs *in vivo*, we compared two independent models for acute, mild stress in mice expressing YFP in CA1 PCs (Thy1-YFP-H): 1) foot shock (FS) and 2) predator odor (PO). Corticosterone levels were mildly elevated in blood plasma 20 minutes after FS and PO acute stress paradigms (Sup. Fig.1a-b). These data fit with the initial phase of the stress response, since plasma cortisol levels have been reported to significantly increase only 30 minutes to an hour after stress induction (46–48). In both paradigms, we found a significant increase in spine density compared to unstressed animals (Fig.1a-c). In addition, acute stress using the FS paradigm shifts spine morphology towards more mature types (Fig.1e,d): mushroom and stubby (49,50). In PO experiments, both the increase in spine density and the shift in spine morphology (Fig.1f) were less prominent compared to FS. Since acute stress-induced changes of corticosterone levels in the hippocampus and other brain regions takes at least 30 minutes, (51,52) a systemic component is very unlikely to be involved in the structural changes in spine density and morphology we find within 20 minutes after acute stress induction.

**Figure 1.**
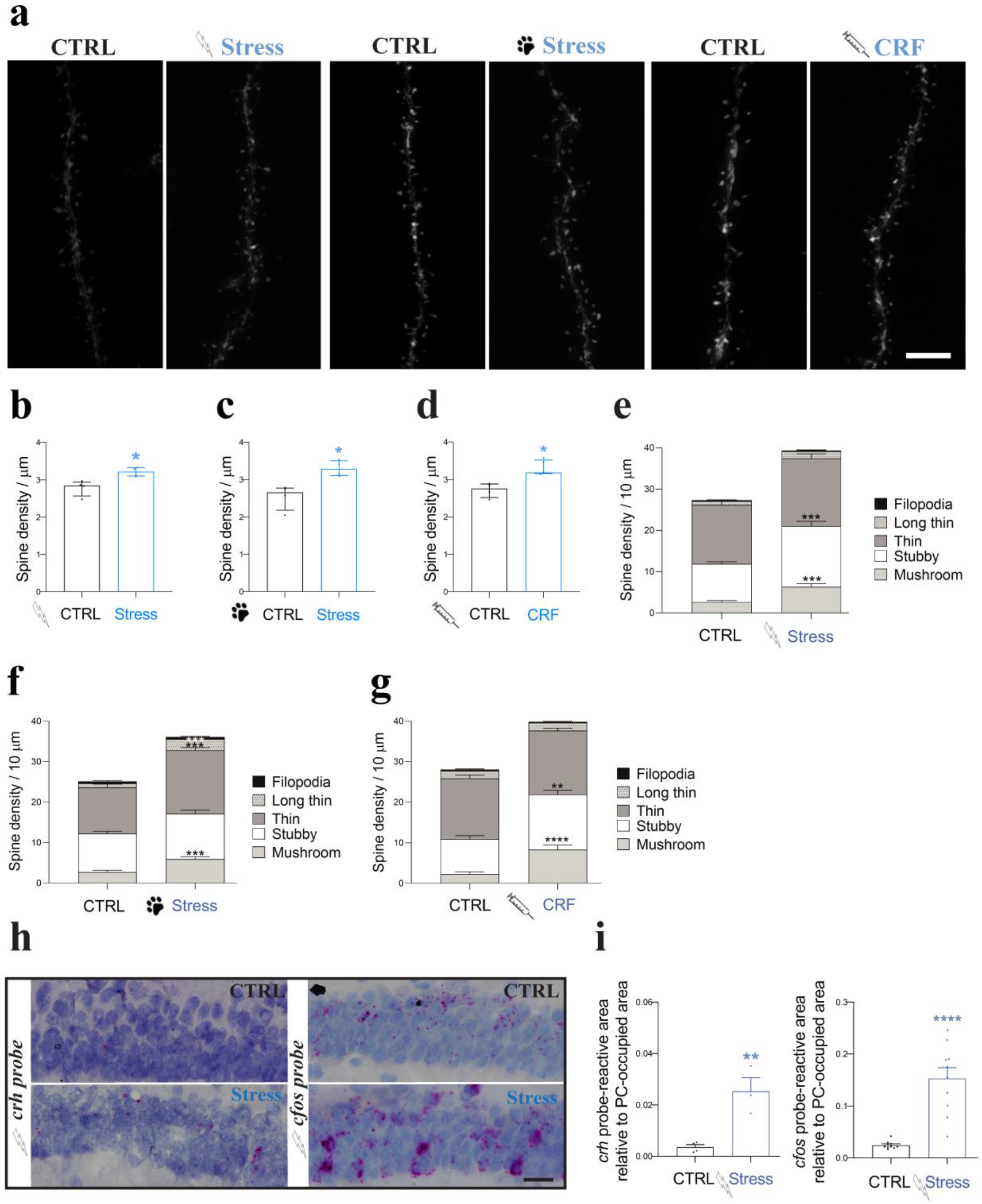
Acute stress and CRF increase spine density in CA1 hippocampus *in vivo*. (a) Representative images of CA1 PC dendrites from Thy1GFP mice before and after acute stress. Left two images: foot shock (FS) paradigm, middle two images: predator odor (PO) paradigm, right two images: 20 minutes 100nM CRF treatment using stereotactic injections into the hippocampal formation. Scale bar=5µm (b-d) Quantification of spine densities after FS, PO and CRF treatment (shown as median with IQR, CTRL: N=4, n=34; FS: N=4, n=34; PO: N=4, n=31; acute CRF treatment: N=5, n=34; Mann-Whitney tested (U=0); * p<0.05). Acute stress paradigms FS (b), PO (c) and acute CRF treatment (d) increase spine density. (e-g) Quantification of spine types. Acute stress paradigms FS (e) (shown as the mean±SEM, CTRL: N=4, n=19; FS: N=5, n=18; multiple t test (filopodia; t ratio=0.6804, long thin; t ratio=2.696, thin; t ratio=1.513, stubby; t ratio=4.728, mushroom; t ratio=4.363). ***p<0.0001). PO (f) (shown as the mean±SEM, CTRL: N=5, n=18; PO: N=4, n=18; multiple t test filopodia; t ratio=0.2124, long thin; t ratio=4.100, thin; t ratio=4.141, stubby; t ratio=0.9460, mushroom; t ratio=4.868). ***p<0.0005) and acute CRF treatment (g) (shown as the mean±SEM, CTRL: N=3, n=14; CRF injections: N=3, n=15; multiple t test filopodia; t ratio=0.8867, long thin; t ratio=0.2817, thin; t ratio=0.9384, stubby; t ratio=3.645, mushroom; t ratio=4.784). **p<0.005) promote spine maturation in PCs CA1. (h) FS increases *crh* (left) and *cfos* (right) mRNA expression in CA1 PCs. Scale bar=25µm (i) Quantification of *crh* (left) and *cfos* (right) mRNA expression (shown as the mean±SEM from >3animals; FS CTRL: N=10 sections; FS: N=10; unpaired t-test (t=2.765, df=18). *p<0.05).

Next, we performed stereotactic injections of CRF into CA1 of YFP-expressing mice to determine whether CRF has the same effect on spines as acute stress. Indeed, CRF significantly increased spine density compared with control (Fig.1a,d) and induced a shift in spine morphology towards more mature types (mushroom and stubby), comparable to the acute stress paradigms (Fig.1e-g).

To confirm our direct stress response in the hippocampus we performed *in situ* hybridization analysis for immediate early genes *crh* and *cfos*. (53–55), in mice 20 minutes after being subjected to FS. We observed a local increase of *crh* and *cfos* mRNA expression in the CA1 PC layer (Fig.1h-i), demonstrating an upregulation of immediate early genes in the CA1 PC layer directly after acute stress.

### *Acute CRF exposure increase*s *the spine density of CA1 pyramidal cells*

To allow a more detailed analysis of the molecular pathway and functional consequences of acute CRF exposure in CA1 PCs, we investigated if the effect of direct CRF injections on spines can be recapitulated in acute hippocampal slices. Indeed, short-term CRF application significantly increased spine density and maturation of dye-filled PCs in acute hippocampal slices (Fig.2a,b).

**Figure 2.**
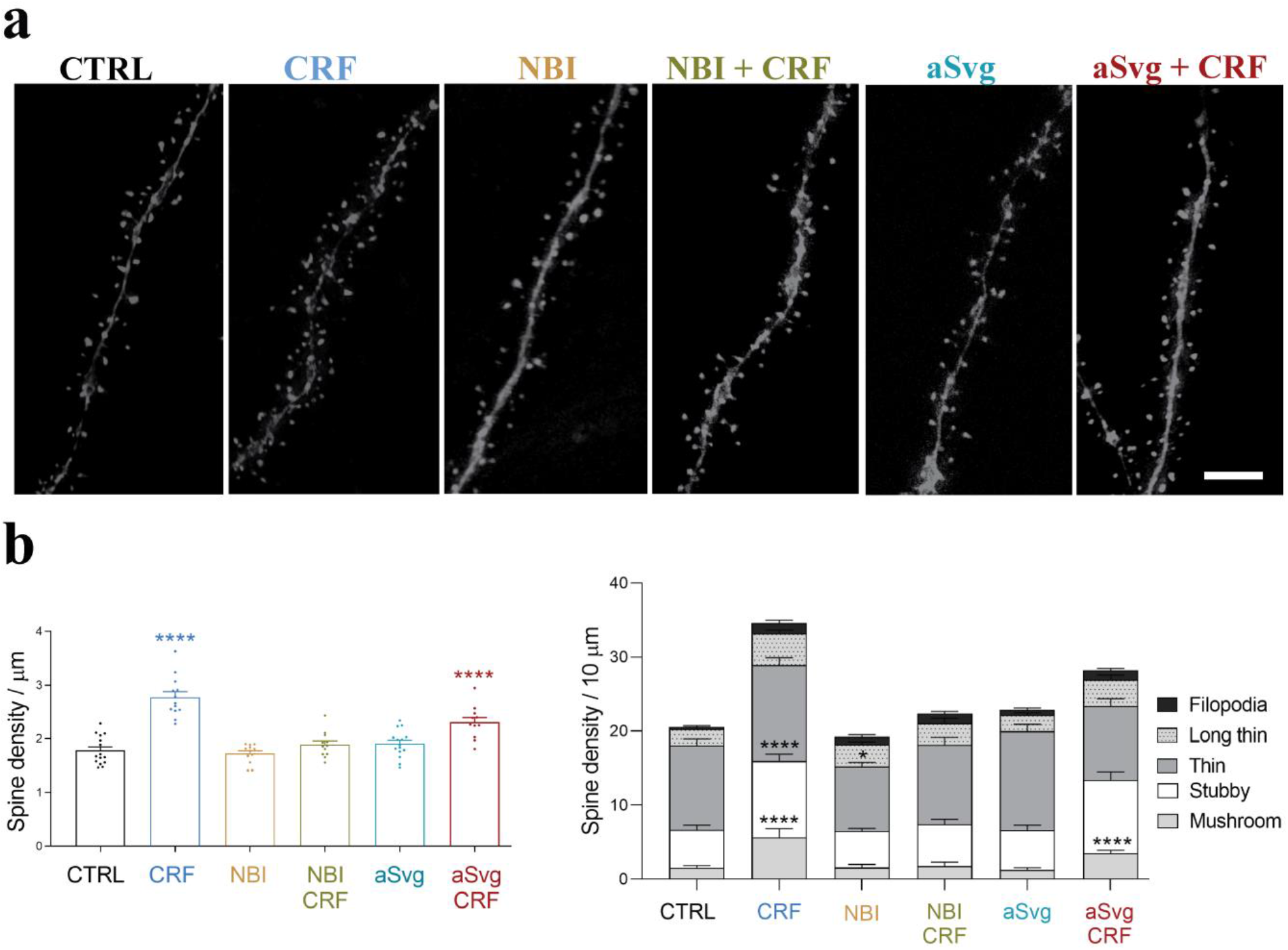
Short-term CRF application increases pyramidal cell spine density and maturation *ex vivo*. (a) Spines on CA1 PC dendrites filled with Alexa 568 using: no treatment (CTRL), only 100nM CRF for 20 minutes, selective CRF-R1 antagonist NBI (1.2μM, NBI), combined NBI and CRF (NBI+CRF), selective CRF-R2 antagonist aSvg (150nM, aSvg) and combined aSvg and CRF (aSvg+CRF) application. Scale bar=5µm. (b-c) Quantification of (b) spine densities (shown as the mean±SEM, CTRL; N=3, n=15; CRF: N=5, n=13; NBI: N=4, n=12; NBI+CRF: N=5, n=12; aSVG: N=5, n=15; aSVG+CRF: N=4, n=9; one-way ANOVA with Dunnett’s multiple comparisons test (F=21.25). *p<0.05, ****p<0.0001), (c) and type (shown as the mean±SEM; two-way ANOVA with multiple comparisons (F=21.25). *p<0.05, ***p<0.005, ****p<0.0001) using aforementioned conditions.

Using acute slices, we set out to identify the underlying CRF receptors involved in mediating the acute spine changes, by pretreating acute slices with their selective antagonists (CRF-R1:NBI 27914 (NBI); K_i_=1.7 nM, CRF-R2: Antisauvagine-30 (aSvg); K_i_=1.4 nM) immediately before CRF treatment. Application of either antagonist alone did not significantly affect spines of CA1 PCs (Fig.2a,b). Inhibition of CRF-R1s completely blocked the CRF-induced increase in spine density and maturation (Fig.2a-c), while inhibition of CRF-R2 partially blocked this CRF effect (but significant, p=0.0027). Together, these data show that the acute CRF-induced increase in mature spine number is predominantly dependent on CRF-R1 signaling, although CRF-R2s can play a complementary role, potentially requiring the deployment of calcium stores (56).

The changes in spine density and maturation sustained at least 1.5 hours after removal/wash out of CRF, suggesting these are long-lasting structural modifications (data not shown).

### Acute CRF exposure modulates functional properties of Schaffer Collateral input into CA1

To determine if the CRF-induced increase in (mature) spine density translates into enhanced functional synaptic connections, we set out to study synaptic function, starting with recording miniature excitatory postsynaptic potentials (mEPSCs) in CA1 PCs. Using pre-treated acute slices with 100nM CRF (incubation started 15 minutes before and continued throughout the recordings), we showed a robust increase in mEPSC frequency, but not amplitude (Fig.3a-e). This finding suggests or an increase in the number of excitatory synapses, or an increase in release probability of individual synaptic connections, or an increase in neuronal network activity. We already found a CRF-induced increase in mature spine density, in line with an increase in functional synaptic connections. However, we also found ultrastructural changes within synapses, that are in line with an increase in release probability (see below) and CRF-induced enhanced network activity as evident from our calcium imaging of CA1 PCs in *ex vivo* acute slices from mice expressing the fluorescent calcium indicator, GCaMP6s (Sup.Fig.2, Sup.Video1). Together, these data suggest that the CRF-induced increase in mEPSC frequency is due to a combination of structural and functional adaptations.

**Figure 3.**
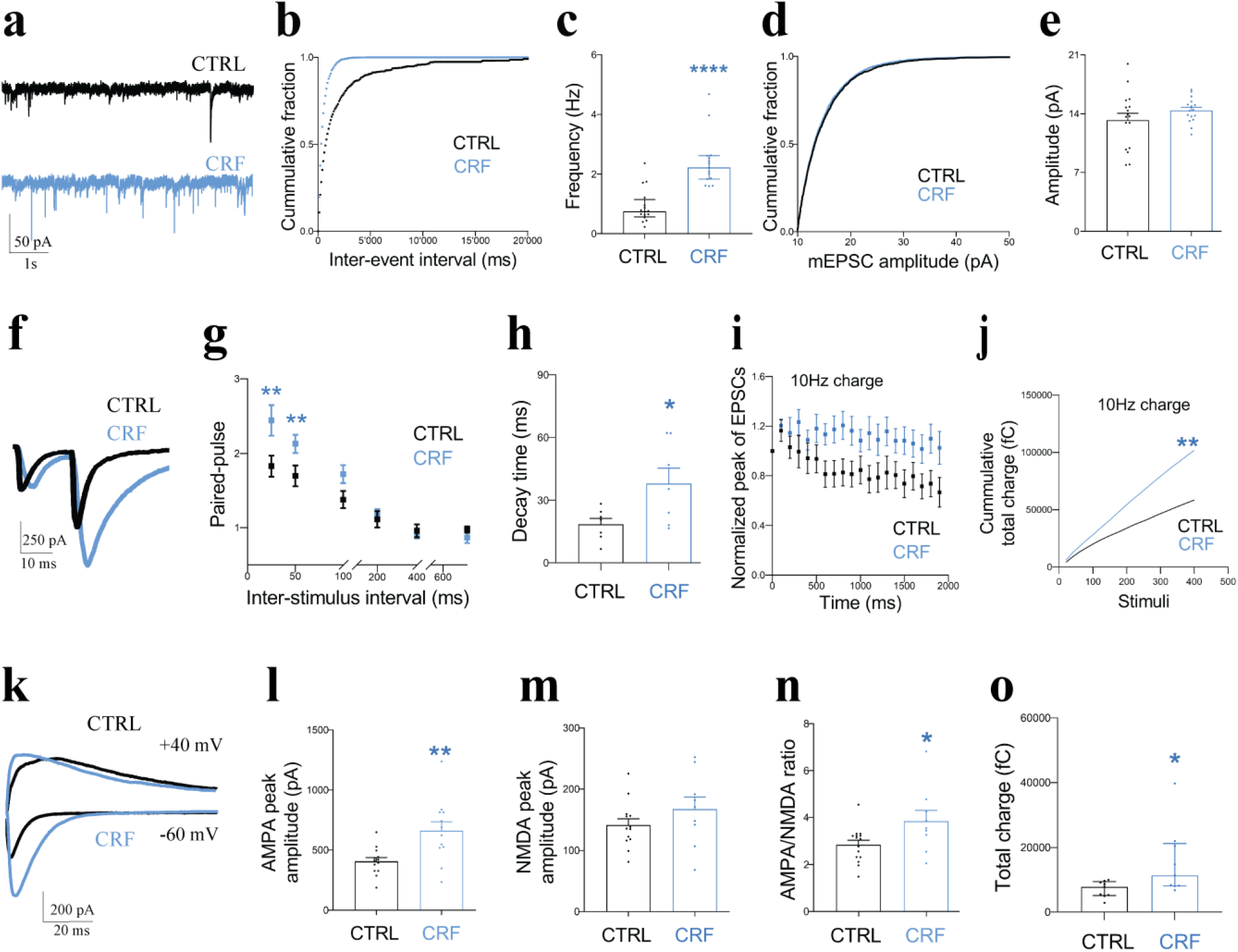
Acute CRF treatment increases synaptic input and synaptic reliability. (a) mEPSCs recorded in CA1 PCs in control (black) and 20 minutes after 100nM CRF treatment (blue). (b-c) CRF increased mEPSCs frequency (shown median with IQR. CTRL: N=3, n=16; CRF: N=3, n=15; Mann-Whitney test (U=15). ****p<0.0001), but not amplitude (d-e) (shown as the mean±SEM. CTRL: N=3, n=17; CRF: N=4, n=16; unpaired t test (t=1.267, df=31). P=0.2147). (f) Stimulation of Schaffer collaterals resulting paired pulse input in recorded CA1 PCs under control (black) and CRF pretreated conditions (blue). (g,h) CRF increased amplitude (increased paired pulse facilitation with 25 and 50 ms inter-stimulation intervals) (g) (shown as the mean±SEM. CTRL: N=4, n=15; CRF: N=4, n=16; unpaired t test (for 25ms; t=3.406 and df=31, for 50ms; t=2.835 and df=31). **p<0.01) and the decay time (h) (shown as mean±SEM. CTRL: N=4, n=15; CRF N=4, n=16; unpaired t test (t=3.738 and df=31). ***p<0.001) of the second evoked EPSC. (i-j) Normalized EPSC amplitude (i) and cumulative total charge released during train stimulation (10 Hz, 200 stimuli) (j) in control (black) and CRF treated (blue) (shown as the median with IQR. CTRL: N=5, n=17; CRF: N=6, n=15; Mann-Whitney test (U=58). **p<0.01). (k) CRF increased AMPAR-mediated EPSC amplitude at SC-CA synapses(Vm=-60mV, black) (l) (shown as the mean±SEM. CTRL: N=4, n=13;CRF: N=3, n=12; unpaired t test (t=3.189, df=23). **p<0.005), but not NMDAR-mediated EPSC amplitude (Vm=+40mV, blue) (m) (mean±SEM. CTRL: N=4, n=13; CRF: N=3, n=9; are unpaired t test (t=1.257, df=20). P=0.2234). (n) AMPAR/NMDAR ratio (shown as mean±SEM. CTRL: N=4, n=15; CRF: N=3, n=9; unpaired t test (t=2.319, df=22). *p<0.05). (o) CRF increases total charge transfer during AMPAR-mediated EPSCs (shown as median with IQR. CTRL: N=4, n=9; CRF: N=3, n=9; Mann-Whitney test (U=17). *p<0.05).

To explore alterations of presynaptic release probability in more detail, we used electrical stimulations of the Schaffer collateral (SC) pathway projecting onto the CA1 PCs. Using paired pulse stimulations, we found increased facilitation (PPF) with 25ms and 50ms intervals in the presence of CRF (Fig.3f-j), suggesting a change in the functional organization of SC presynaptic terminals. We observed a striking increase in the decay time constant in CRF-treated slices (Fig.3h), suggesting increased sustained/asynchronous release following evoked release. To further explore the effect of acute CRF exposure during more demanding periods of SC input activity, we performed train stimulations and analyzed both the synchronous peak amplitude and the total cumulative evoked charge. We observed decreased synaptic fatigue during 10Hz train stimulations (Fig.3i) and an almost 2-fold increase in the absolute total cumulative charge after CRF treatment (Fig.3j). Together, these observations suggest that CRF changes presynaptic function, ultimately resulting in enhanced synaptic reliability.

To determine if CRF indeed affects the number of mature/functional synaptic contacts (as suggested by mEPSC frequency and changes in spines), we stimulated SC inputs and consecutively recorded AMPA- and NMDA-receptor mediated evoked EPSCs (−60 mV and +40 mV respectively, Fig.3k-n). CRF-treatment induced a significant increase in AMPA component, both amplitude and total charge (Fig.3l, o), while NMDA amplitude was unaltered. Consequently, CRF increased the AMPA/NMDA ratio suggesting a shift towards mature/functional synaptic connections, in line with our spine analysis data.

To explore the long-term effects of CRF on synaptic plasticity and network function, we examined long-term depression (LTD) and long-term potentiation (LTP) of the SC pathway onto CA1 PCs, using multi-electrode array (MEA) extracellular field potential recordings (field excitatory postsynaptic potentials, fEPSPs) (Fig.4). In the cerebellum, LTD induction requires CRF (57), but this CRF-dependency of LTD generation has not been reported in the hippocampus (58–60). First, we confirmed that - in our hands - we were able to induce substantial LTD and subsequently if this plasticity paradigm was reversible, using a consecutive LTP induction on the same acute slices (Fig.4a). Next, we investigated the effect of acute CRF application on baseline fEPSP amplitude and subsequently on either LTD or LTP in separate experiments (Fig.4b-e). During CRF application (15 minutes, indicated with “15’ CRF” (Fig.4b,e,h) we observed a clear increase in fEPSP amplitude, likely representing the short-term increase in synaptic function/reliability described above. This increase in fEPSP was transient and after CRF wash out, the amplitude returned to baseline, as previously described (27). Intriguingly, CRF treatment significantly enhanced LTD and LTP induction (Fig.4c,e), seemingly increasing the spectrum and/or sensitivity of long-term plasticity mechanisms. To determine the involvement of CRF-Rs in enhancing LTP, we combined application of the CRF-receptor antagonists NBI and aSvg with LTP induction. By themselves, these blockers did not affect baseline fEPSP amplitudes or LTP induction (Fig.4f). Combined with CRF treatment, blocking either of the two CRF-Rs did not inhibit CRF-induced enhancement of LTP (Fig.4h,i). However, combining both blockers abolished the acute CRF-dependent LTP enhancement, indicating that activation of either receptor is sufficient for this form of plasticity.

**Figure 4.**
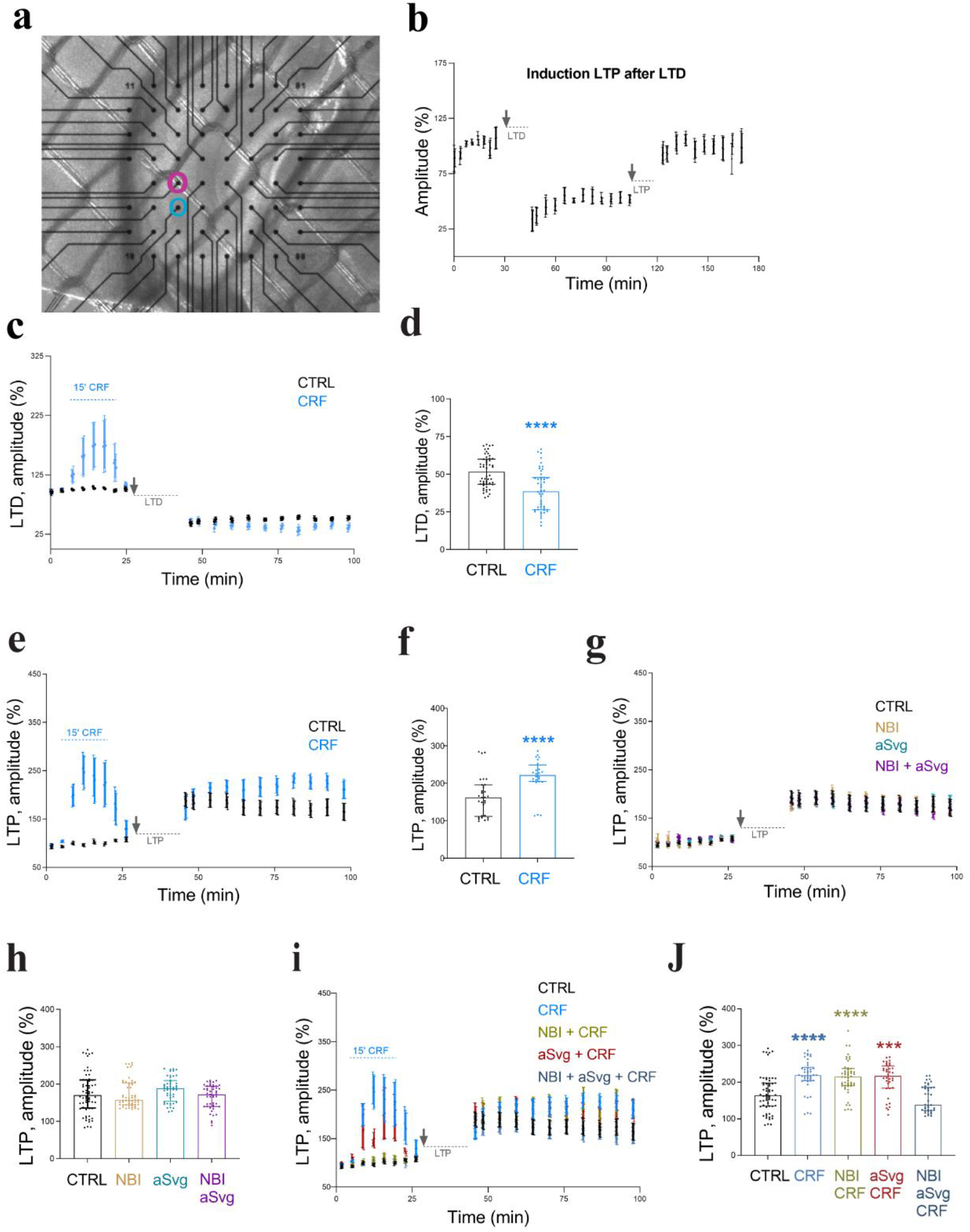
CRF can augment long-term synaptic plasticity via CRF-R1 or CRF-R2 activation. Multi electrode array recording of evoked fEPSP from the SC in the CA1. (a) Image of a mouse acute hippocampal slice on the multi electrode array (MEA2100; Multichannel Systems) used for field excitatory post-synaptic potential (fEPSP) recordings with the stimulation electrode (blue) and recording electrode (pink) to stimulate Shaffer collateral-CA1 connections. (b) Consecutive long-term depression (LTD) and long-term potentiation (LTP) induction. Baseline fEPSPs were recorded for approximately 25 minutes, LTP induction protocol was applied to the same slices and recording continued for another 60 minutes. (c) –LTD in control slices (black) and slices treated with CRF (15 minutes, 100nM CRF). Treatment period indicated with dashed line (blue)). (d) Averaged fEPSC amplitude 60 minutes after LTD induction (normalized to baseline) (shown as the median with IQR. CTRL: N=9; CRF: N=8; Mann-Whitney test (U=631). ****p<0.0001). CRF treatment increased LTD by 17% compared to control (e) – LTP in control slices (black) and slices treated with CRF (15 minutes, 100nM CRF. Treatment period indicated with dashed line (blue)). (f) CRF increased LTP efficiency by 32% (shown as the median with IQR. CTRL: N=11; CRF: N=9; Mann-Whitney test (U=139). ****p<0.0001). (g) LTP induction in combination with either CRF-R1 blocker (NBI 1.2µM), CRF-R2 blocker (aSvg 150nM) or both. (h) CRF-R blockers do not affect LTP (shown as the median with IQR. CTRL: N=11; NBI: N=9; aSVG: N=8; NBI+aSVG: N=8; Kruskal-Wallis test with Dunn’s multiple comparisons test (Kruskal-Wallis statistic=6.357)). (i) LTP induction using combinations of CRF-Rs blockers. Blockers were present throughout the recording. (j) Effect of CRF on LTP can be established via both CRF-Rs pathway, but no additivity was found if both pathways are available (shown as the median with IQR. CTRL: N=11; CRF: N=9; NBI: N=9; NBI+CRF: N=9; aSVG: N=8; aSVG+CRF: N=8; NBI+aSVG: N=8; NBI+aSVG+CRF: N=8; Kruskal-Wallis test with Dunn’s multiple comparisons test (Kruskal-Wallis statistic=94.32). ***p<0.0005, ****p<0.0001).

### Acute CRF exposure leads to ultrastructural alterations of synapses

To further scrutinize the short-term effects of CRF on synaptic structure and organization, we performed ultrastructural electron microscopy (EM) analysis on hippocampal *ex vivo* acute slices, focusing on synaptic connections on CA1 PCs in the stratum radiatum (SR), the layer Schaffer collateral synapses are predominantly located Fig.5, Sup.Video2,3).

**Figure 5.**
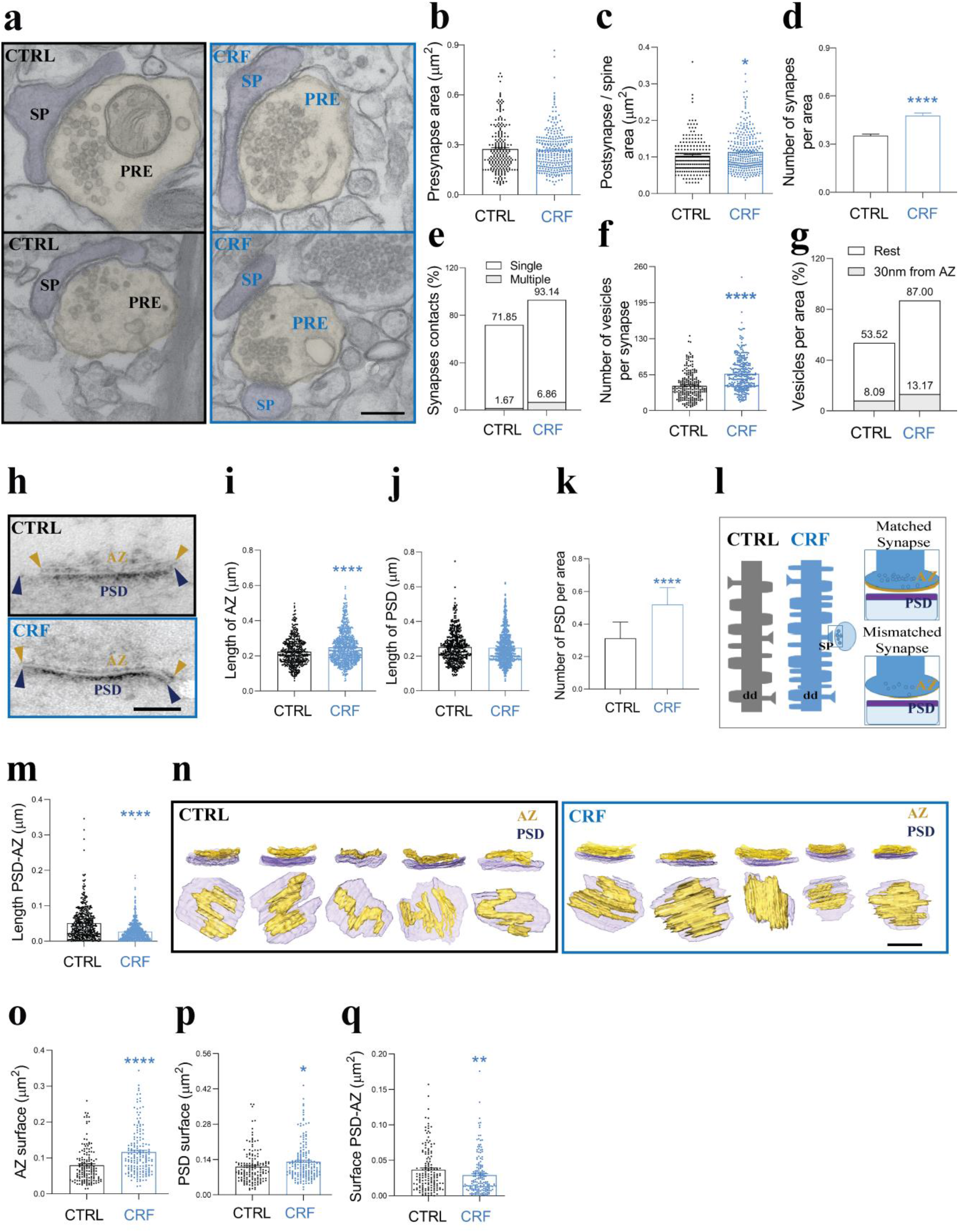
Acute CRF alters multiple aspects of synaptic architecture. (a) TEM images from control (CTRL) and CRF-treated acute slices. Scale bar=200nm. (b,c) Quantification of presynaptic (PRE), (b) (shown as the median with IQR. CTRL: N=3, n=216; CRF: N=3, n=283; Mann-Whitney test (U=30100). P=0.7715) postsynaptic (c) areas (spines, SP) (shown as the median with IQR. CTRL: N=3, n=219; CRF: N=3, n=304; Mann-Whitney test (U=29456). *p<0.05). (d,e) CRF (100nM, 20 minutes) increased both the number of synapses per area (d) (shown as the median with IQR. CTRL: N=3, n=177; CRF: N=3, n=181; Mann-Whitney test (U=10819). ****p<0.0001) and the number of multiple postsynaptic boutons per single presynapse (e). (f,g) CRF increased the total number of vesicles per synapse (f) (shown as the median with IQR. CTRL: N=3, n=220; CRF: N=3, n=305; Mann-Whitney test (U=18748). ****p<0.0001), and reorganized synaptic vesicles towards the active zone (AZ) (g). (h) TEM images of PTA-stained synapses. Scale bar=100nm. (i-k) CRF increased AZ length (i) (shown as the median with IQR. CTRL: N=3, n=452; CRF: N=3, n=771; Mann-Whitney test (U=147985). ****p<0.0001) and the number of PSD per area (k) (shown as the median with IQR. CTRL: N=3, n=150;) CRF: N=3, n=150; Mann-Whitney test (U=3736). ****p<0.0001), but not postsynaptic density (PSD) length (j) (shown as the median with IQR. CTRL: N=3, n=452; CRF: N=3, n=771; Mann-Whitney test (U=167545). P=0.2610) (l) Relationships between AZ and PSD at the synapse. The upper right represents an idealized, matched synapse, where the lengths of AZ and PSD are approximately equal in length, while the lower a mismatched synapse where the length of the PSD is (typically) larger than the AZ. (m) CRF enhanced synaptic matching between AZ and PSD (shown as the median with IQR. CTRL: N=3, n=452; CRF: N=3, n=771; Mann-Whitney test (U=109620). ****p<0.0001). (n) 3D reconstruction of AZ and PSD after FIB-SEM imaging shows a CFR dependent increase in AZ surface area (yellow). Scale bar=150nm. (o-q) Quantification of AZ-PSD complexity using 3D reconstructed images, confirms CRF increased AZ surface area (o) (shown as the median with IQR. CTRL: N=3, n=150; CRF: N=3, n=173; Mann-Whitney test (U=8026). ****p<0.0001), increased PSD surface area (p) (shown as the median with IQR. CTRL: N=3, n=150; CRF: N=3, n=173; Mann-Whitney test (U=10922). *p<0.05) and promoted tighter matching between them (q) (shown as the median with IQR. CTRL: N=3, n=150; CRF: N=3, n=173; Mann-Whitney test (U=10737). **p<0.01).

CRF did not affect presynaptic bouton area (Fig.5a,b). However, we did observe an increase in postsynaptic compartment size (Fig.5c), the number of synapses per area unit (Fig.5d), and number of single presynaptic terminals innervating multiple spines (Fig.5e). To investigate whether CRF induces structural changes within presynaptic terminals, we analyzed the number and localization of synaptic vesicles. Indeed, CRF increases the total number of vesicles per synapse area and in addition repolarizes these vesicles towards the release sites, resulting in more vesicles within 30 nm from active zone (AZ))(Fig.5f,g).

To evaluate the spatial relationship between AZ and post-synaptic density (PSD), we stained slices with PTA, which highlights macromolecular complexes of AZ/PSD in the synapse (61,62). We focused on asymmetric synapses at the CA1-SR, where secondary and tertiary dendrites of PCs are located and SC synapses are predominantly located (Fig.5h-k). CRF induced a significant increase in the number PSDs (corrected for postsynaptic terminal area) and in the length of AZ, without alterations in PSD length. These findings prompted us to investigate the alignment between the AZ and PSD (Fig.5l). In a “matching” synapse, the size of the AZ and PSD are comparable to each other (Fig.5l, top), whereas in “mismatched” synapses the AZ is smaller (Fig.5l, bottom). Using this approach, we found CRF significantly increased matching between AZ and PSD length (Fig.5m).To further examine synapse matching, we utilized FIB-SEM based imaging, to allow a more detailed three-dimensional analysis of synapse ultrastructure (Fig.5n-q, Sup.Video2,3). The 3D-reconstructed spatial organization of AZ-PSD complexes confirmed a CRF-induced increase in AZ surface but also showed the previously missed increase in PSD size. In addition, we confirmed that CRF signaling facilitated AZ-PSD matching (Fig.5n,q).

## Discussion

Acute stress has a diverse range of beneficial effects on brain function (63,64) and multiple studies have demonstrated the involvement of CRF as a central regulator in this adaptive process (12,65– 68). However, the acute role of CRF as a local neuromodulator in structural and functional synaptic plasticity has not been investigated extensively. Here, we provide detailed insights into the acute role of CRF as a local neuropeptide in acute stress. Our research shows that the structural and functional consequences of acute stress paradigms can be recapitulated both *in vivo and ex vivo*, using short-term application of physiologically relevant CRF concentrations (8)

CRF treatment (injected *in vivo* or applied to acute slices *ex vivo*) resulted in similar structural adaptations as observed during acute stress paradigms, suggesting a prominent role of CRF in regulating physiological responses to acute stress. Short-term CRF treatment resulted in rapid structural and functional adaptations, leading to an overall increase in functional synaptic contacts. In short, we showed that CRF 1) increased spine density and maturation, 2) increased synapse number and size, 3) revised synaptic vesicle organization towards release sites, 4) enhanced matching of synaptic contact, 5) increased synaptic efficacy and 6) enhanced the functional range of long-term plasticity. Systemically released stress hormones likely cannot be involved in the direct effects that we found after acute stress and CRF application, since these have been reported to reach brain tissue well after the structural and functional changes we describe here (51,52). However, there probably is a temporal integration of immediate (initiated by the local release of neuromodulators) and delayed (through systemically derived hormones) stress responses within brain regions. Our *in vivo* data showed an upregulation of immediate early gene *crh* and *cfos mRNA* expression (Fig.1i,j) after acute stress, indicating responses likely also involve widespread long-term changes in neuronal function. In addition, our *ex vivo* results confirm this local response by treatment of CRF and absence of hormonal response. Together, our findings indicate a prominent role of locally released CRF during the immediate phase of acute stress, modulating synaptic input in the CA1 PCs.

Activation of CRF-R1 is required for CRF-induced changes in spine density and maturation, while CRF-R2s are not (Fig.2b,c). Since CRF-R1 activation is also a prerequisite for inducing *cfos* expression (69,70), the observed structural changes might depend on processes downstream of *cfos* signaling. Comparably, the transient increase in fEPSP amplitude during CRF applications requires CRF-R1 activation. Since the observed CRF-R1-dependent structural changes would presumably persist after CRF exposure, it seems more likely that the reversible CRF-R1 dependent increase in fEPSP responses is due to a transient increase in presynaptic efficacy via CRF-R1s expressed in the presynaptic compartment (71,72). Indeed, both evoked and spontaneous synaptic input in CA1 PCs increased while applying CRF indicating an immediate effect of CRF on synapse function. In contrast, either CRF-R1 or CRF-R2 activation (or both) was sufficient to enhance long-term plasticity (Fig.4h,-i). Our data supports CRF as a positive regulator of synaptic transmission, in agreement with other reports describing CRF generally as a facilitator of excitatory neurotransmission throughout different brain regions (31,55,73,74).

PPF is an activity dependent increase in pre-synaptic release probability due to accumulation of presynaptic Ca^2+^. CRF increases paired pulse facilitation (PPF) of the Schaffer Collateral synapses (Fig.3g), likely due to the relocation of synaptic vesicles towards the active zone (Fig.5g) which would increase either the size or the replenishment rate of the release pool. The CRF-induced increase in synaptic vesicle number and redistribution towards the active zone is expected to also affect synaptic release efficacy during sustained periods of activity, which is indeed what we observed during train stimulations (Fig.3i). EM confirmed the increase in the docking pool of vesicles by CRF (Fig.5f,g), thereby providing evidence of the mechanism of action of CRF in acute stress response by enhancing structural architecture and functional properties of the synaptic network.

In contrast to CRF-R1, there is still much debate over the presence of CRF-R2 in the rodent hippocampus (7,10,75–79). Some reports confirm expression of CRF-R2 (71,75,76,78), while others disregard its presence (9,80,81). CRF-R2 mRNA has been reported throughout the hippocampal formation, albeit in lower amounts compared to CRF-R1 (71,75). Potentially the presence of different isoforms of CRF-R2 (full-length and truncated) underlies these contradicting reports (36). In addition, the two receptors are also known to have different kinetics. While CRF-R1 is activated fast in acute stages, studies in knock-out mice suggest that CRF signaling via CRF-R2 has a slower kinetic (4,82,83). Indeed, CRF-receptors were insufficient to establish the acute CRF-induced enhancement of synaptic efficacy (as measured by a transient increase in fEPSPs) but consecutive LTP induction was enhanced, in line with a delayed role of CRF-R2 in acute CRF signaling.

In conclusion, we report that acute CRF signaling in CA1 PCs involves a complicated interplay of morphological and functional synaptic adaptations, which culminate in enhancing both short- and long-term responsiveness of the underlying neuronal network, potentially affecting hippocampus dependent learning strategies during short stressful events.

## Acknowledgements

We thank Drs. Matthew Holt, Linette Lim, and Bart De Strooper (VIB-KU Leuven) for critical reading of the manuscript, VIB BioImaging Core – Leuven and Ghent for help with imaging, Eline Creemers for help with electrophysiology. We thank Drs. Karl Farrow, Vincent Bonin (NERF) for the Thy1-GCaMP6s mouse line, the behavioral core (mINT-KU Leuven) Dr.Zsuzsanna Callaerts-Vegh. We thank Dr. Katarzyna Marta Zoltowska and Dr. Lucía Chávez-Gutiérrez for assistance with the ELISA experiment. DV is supported by a Methusalem grant from KU Leuven and the Flemish Government awarded to Dr. Bart De Strooper (METH/14/07). FWO, Image Storage Platform for Analysis Management and Mining project (ISPAMM; AKUL/13/39).

## Author contributions

N.G., D.V., and V.R. conceived the project and N.G. and D.V. designed the experiments. D.V., K.V., P.B. and K.Z. performed experiments. D.V., N.G. and K.V. analyzed the data. K.W. designed experiments involving electrophysiology, contributed reagents, materials, and analysis tools. L.D.G. and L.M. provided the Thy1-YFP-H mouse line. N.G., D.V. and K.W. wrote the paper.

## Supplemental Figures

**Supplementary Figure 1.**
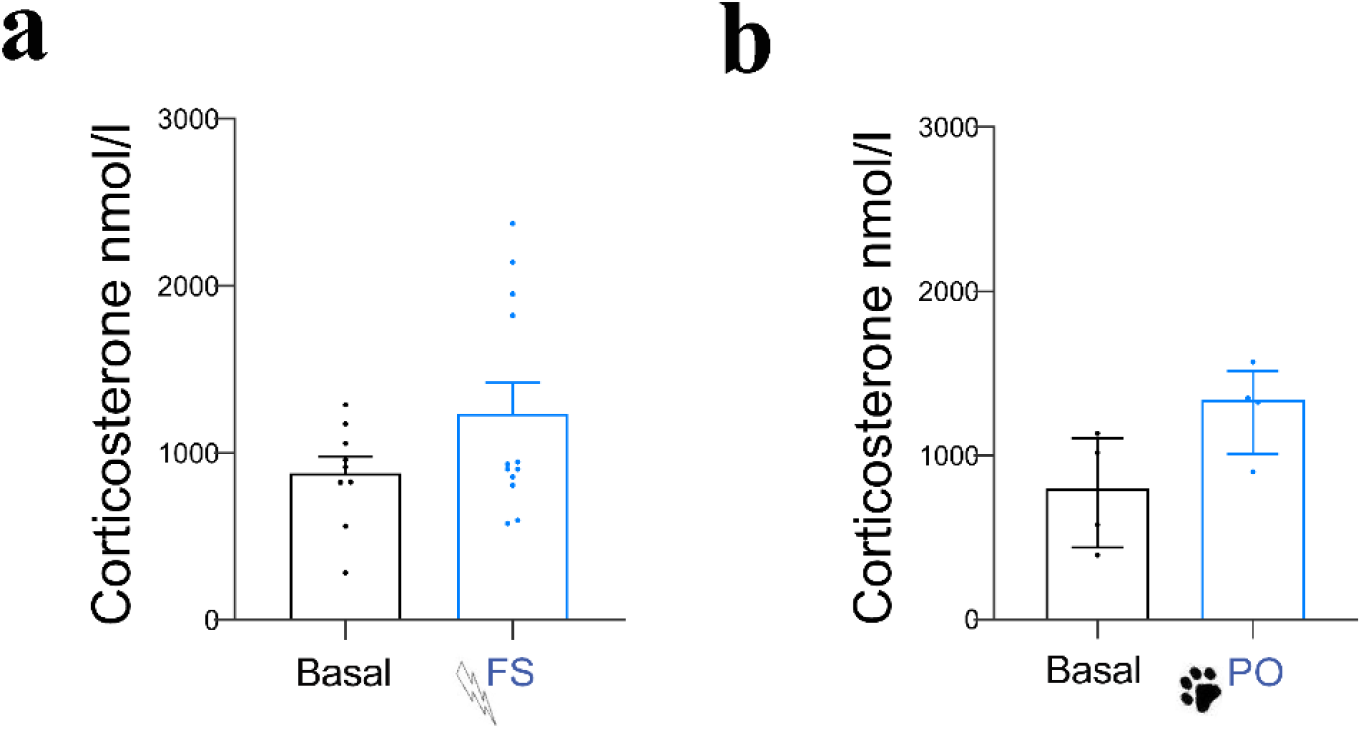
Blood plasma levels in acute stress paradigms. (a) Quantification of corticosterone (CORT) concentrations in blood plasma after FS (a) (shown as mean±SEM, CTRL: N=9; FS: N=12; unpaired t-test (t=1535, df=19). p=0.1413) and PO (b) (shown as median±IQR, CTRL: N=4; PO: N=4; Mann-Whitney test (U=2). p=0.1143).

**Supplementary Figure 2.**
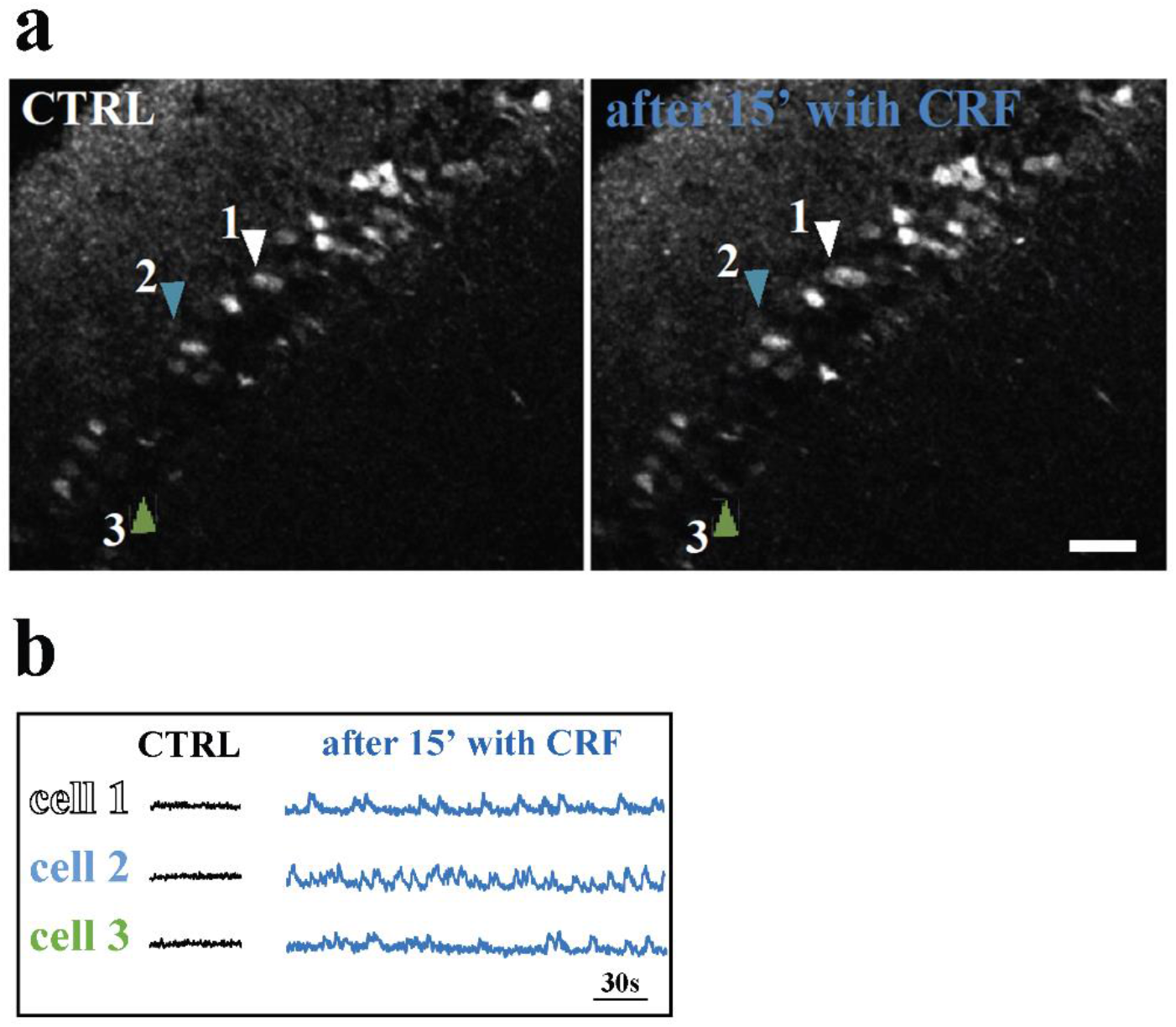
Acute CRF exposure increases calcium release *in vivo* in CA1 PCs. (a) Image of GCAMP+ PCs CA1 before treatment with CRF (left) and same field of view after 15 minutes with CRF 100 nM (right). Arrows indicate same cells in control and CRF conditions. Scale bar 50µm. (b) the traces of calcium influx of individual cells marked with arrows as in a, before (black) and after CRF application (blue).

## Supplementary Video Legends

**Supplemental Video 1**.

**Acute CRF exposure increases calcium release in CA1 PC layer *in vivo*, in slices prepared from mice expressing the green fluorescent calcium indicator, GCaMP6 (Thy1-GCaMP6 mice)**. (Left) two-photon microscope imaging using 20x objective started in aCSF, with a capture of 300 images (average of 15 frames per image), with a 30 ms interval. After 300 images, slice was continue perfused with 100 nM CRF in aCSF, and other 300 images were taken. (Right) after 15 minutes with CRF in aCSF, the same field of view was imaged last time, 600 images were taken with the same settings. Arrows indicate same cells in control and CRF condition. Abbreviations: aCSF-artificial cerebrospinal fluid, Ca-calcium, CTRL - control, CRF - corticotropin-releasing factor.

**Supplemental Video 2**.

**FIB-SEM imaging and reconstruction of PTA staining in control hippocampal slices**. The FIB-SEM was set to remove 5-nm-thick layers and image acquisition was done using a backscattered electron detector at 1.5 kV (0.005 µm/pixel), at 5kX magnification. Individual segmentation of an AZ shown in yellow and PSD segment in purple (N=1). Abbreviations: AZ - active zone, CTRL - control, CRF - corticotropin-releasing factor, PSD - postsynaptic density, PTA - phosphotungstic acid.

**Supplemental Video 3**.

**FIB-SEM imaging and reconstruction of PTA staining in CRF treated hippocampal slices**. The FIB-SEM was set to remove 5-nm-thick layers and image acquisition was done using a backscattered electron detector at 1.5 kV (0.005 µm/pixel), at 5kX magnification. Individual segmentation of an AZ shown in yellow and PSD segment in purple (N=1). Abbreviations: AZ - active zone, CTRL - control, CRF - corticotropin-releasing factor, PSD - postsynaptic density, PTA - phosphotungstic acid.

## References

1. Selye H. Streß-syndrome. A syndrome produced by diverse nocuos agents. Nature. 1936;

2. Selye H. Stress and the general adaptation syndrome. Br Med J. 1950;

3. Selye H. Confusion and controversy in the stress field. J Human Stress. 1975;

4. Joëls M, Baram TZ. The neuro-symphony of stress. Nat Rev Neurosci. 2009 Jun 2;10(6):459–66.

5. Mcewen BS, Gianaros PJ. Central role of the brain in stress and adaptation: Links to socioeconomic status, health, and disease. Ann N Y Acad Sci. 2010;1186:190–222.

6. Chrousos GP. Stress and disorders of the stress system. Vol. 5, Nature Reviews Endocrinology. 2009.

7. Maras PM, Baram TZ. Sculpting the hippocampus from within: Stress, spines, and CRH. Trends Neurosci. 2012;35(5):315–24.

8. Deussing JM, Chen AA. The Corticotropin-Releasing Factor Family: Physiology of the Stress Response. Physiol Rev. 2018;98:2225–86.

9. Henckens MJAG, Deussing JM, Chen A. Region-specific roles of the corticotropin-releasing factor-urocortin system in stress. Nat Rev Neurosci. 2016;17(10):636–51.

10. Dedic N, Deussing AC and JM. The CRF Family of Neuropeptides and their Receptors - Mediators of the Central Stress Response. Curr Mol Pharmacol. 2017;10:1–28.

11. Vandael D, Gounko NVNV. Corticotropin releasing factor-binding protein (CRF-BP) as a potential new therapeutic target in Alzheimer’s disease and stress disorders. Transl Psychiatry. 2019 Oct 22;9(1):272.

12. Chen Y, Andres AL, Frotscher M, Baram TZ. Tuning synaptic transmission in the hippocampus by stress: the CRH system. Front Cell Neurosci. 2012;6.

13. Gunn BG, Sanchez GA, Lynch G, Baram TZ, Chen Y. Hyper-diversity of CRH interneurons in mouse hippocampus. Brain Struct Funct. 2019;224(2):583–98.

14. Chen Y, Brunson KL, Müller MB, Cariaga W, Baram TZ. Immunocytochemical distribution of corticotropin-releasing hormone receptor type-1 (CRF1)-like immunoreactivity in the mouse brain: Light microscopy analysis using an antibody directed against the C-terminus. J Comp Neurol. 2000;

15. Refojo D, Echenique C, Müller MB, Reul JMHM, Deussing JM, Wurst W, et al. Corticotropin-releasing hormone activates ERK1/2 MAPK in specific brain areas. Proc Natl Acad Sci U S A. 2005;

16. McEwen BS, Gianaros PJ. Stress- and Allostasis-Induced Brain Plasticity. Annu Rev Med. 2010;62(1):431–45.

17. Zoladz PR, Diamond DM. Linear and non-linear dose-response functions reveal a hormetic relationship between stress and learning. Dose-Response. 2009;7(2):132–48.

18. Schwabe L, Joëls M, Roozendaal B, Wolf OT, Oitzl MS. Stress effects on memory: An update and integration. Neurosci Biobehav Rev. 2012;36(7):1740–9.

19. Krugers HJ, Lucassen PJ, Karst H, Joëls M. Chronic stress effects on hippocampal structure and synaptic function: Relevance for depression and normalization by anti-glucocorticoid treatment. Front Synaptic Neurosci. 2010;(JUL).

20. De Quervain DJF, Roozendaal B, McGaugh JL. Stress and glucocorticoids impair retrieval of long-term spatial memory. Nature. 1998;

21. Justice NJ. The relationship between stress and Alzheimer’s disease. Neurobiol Stress. 2018;8:127–33.

22. K. B, K. S, M.-È. T. Chronic stress as a risk factor for Alzheimer’s disease: Roles of microglia-mediated synaptic remodeling, inflammation, and oxidative stress. Neurobiol Stress. 2018;9(February):9–21.

23. Gounko N V., Swinny JD, Kalicharan D, Jafari S, Corteen N, Seifi M, et al. Corticotropin-releasing factor and urocortin regulate spine and synapse formation: Structural basis for stress-induced neuronal remodeling and pathology. Mol Psychiatry. 2013;18(1):86–92.

24. Chen Y, Kramár EA, Chen LY, Babayan AH, Andres AL, Gall CM, et al. Impairment of synaptic plasticity by the stress mediator CRH involves selective destruction of thin dendritic spines via RhoA signaling. Mol Psychiatry. 2013;18(4):485–96.

25. Joëls M, Fernandez G, Roozendaal B. Stress and emotional memory: A matter of timing. Trends Cogn Sci. 2011;15(6):280–8.

26. Swinny JD, Metzger F, Ijkema-Paassen J, Gounko N V., Gramsbergen A, Van Der Want JJL. Corticotropin-releasing factor and urocortin differentially modulate rat Purkinje cell dendritic outgrowth and differentiation in vitro. Eur J Neurosci. 2004;19(7):1749–58.

27. Blank T, Nijholt I, Eckart K, Spiess J. Priming of Long-Term Potentiation in Mouse Hippocampus by Corticotropin-Releasing Factor and Acute Stress: Implications for Hippocampus-Dependent Learning. J Neurosci. 2002;22(9):3788–94.

28. Rebaudo R, Melani R, Balestrino M, Izvarina N. Electrophysiological effects of sustained delivery of CRF and its receptor agonists in hippocampal slices. Brain Res. 2001;

29. Chen Y, Rex CS, Rice CJ, Dubé CM, Gall CM, Lynch G, et al. Correlated memory defects and hippocampal dendritic spine loss after acute stress involve corticotropin-releasing hormone signaling. Proc Natl Acad Sci U S A. 2010;107(29):13123–8.

30. Chen Y, Bender RA, Brunson KL, Pomper JK, Grigoriadis DE, Wurst W, et al. Modulation of dendritic differentiation by corticotropin-releasing factor in the developing hippocampus. Proc Natl Acad Sci U S A. 2004;

31. Aldenhoff JB, Gruol DL, Rivier J, Vale W, Siggins GR. Corticotropin releasing factor decreases postburst hyperpolarizations and excites hippocampal neurons. Science (80-). 1983;221(4613):875–7.

32. Haug T, Storm JF. Protein kinase A mediates the modulation of the slow Ca2+-dependent K+ current, I(sAHP), by the neuropeptides CRF, VIP, and CGRP in hippocampal pyramidal neurons. J Neurophysiol. 2000;

33. Bali A, Jaggi AS. Electric foot shock stress: A useful tool in neuropsychiatric studies. Rev Neurosci. 2015;26(6):655–77.

34. Clark SM, Sand J, Francis TC, Nagaraju A, Michael KC, Keegan AD, et al. Immune status influences fear and anxiety responses in mice after acute stress exposure. Brain Behav Immun. 2014;38:192–201.

35. Wu YP, Gao HY, Ouyang SH, Kurihara H, He RR, Li YF. Predator stress-induced depression is associated with inhibition of hippocampal neurogenesis in adult male mice. Neural Regen Res. 2019;

36. Sterley TL, Baimoukhametova D, Füzesi T, Zurek AA, Daviu N, Rasiah NP, et al. Social transmission and buffering of synaptic changes after stress. Nat Neurosci. 2018;21(3):393–403.

37. Wilcock DM, DiCarlo G, Henderson D, Jackson J, Clarke K, Ugen KE, et al. Intracranially administered anti-Aß antibodies reduce ß-amyloid deposition by mechanisms both independent of and associated with microglial activation. J Neurosci. 2003;

38. Belevich I, Joensuu M, Kumar D, Vihinen H, Jokitalo E. Microscopy Image Browser: A Platform for Segmentation and Analysis of Multidimensional Datasets. PLoS Biol. 2016;14(1).

39. Condomitti G, Wierda KD, Schroeder A, Rubio SE, Vennekens KM, Orlandi C, et al. An Input-Specific Orphan Receptor GPR158-HSPG Interaction Organizes Hippocampal Mossy Fiber-CA3 Synapses. Neuron. 2018;100(1):201–215.e9.

40. Pastoll H, White M, Nolan M. Preparation of parasagittal slices for the investigation of dorsal-ventral organization of the rodent medial entorhinal cortex. J Vis Exp. 2012;(61).

41. Largo-Barrientos P, Apóstolo N, Creemers E, Callaerts-Vegh Z, Swerts J, Davies C, et al. Lowering Synaptogyrin-3 expression rescues Tau-induced memory defects and synaptic loss in the presence of microglial activation. Neuron. 2021;

42. Bessa JM, Ferreira D, Melo I, Marques F, Cerqueira JJ, Palha JA, et al. The mood-improving actions of antidepressants do not depend on neurogenesis but are associated with neuronal remodeling. Mol Psychiatry. 2009;14(8):764–73.

43. Leuner B, Shors TJ. Stress, anxiety, and dendritic spines: What are the connections? Neuroscience. 2013;251:108–19.

44. Sandi C, Davies HA, Cordero MI, Rodriguez JJ, Popov VI, Stewart MG. Rapid reversal of stress induced loss of synapses in CA3 of rat hippocampus following water maze training. Eur J Neurosci. 2003;17(11):2447–56.

45. Magariños AM, García Verdugo JM, Mcewen BS. Chronic stress alters synaptic terminal structure in hippocampus. Proc Natl Acad Sci U S A. 1997;94(25):14002–8.

46. Gong S, Miao YL, Jiao GZ, Sun MJ, Li H, Lin J, et al. Dynamics and correlation of serum cortisol and corticosterone under different physiological or stressful conditions in mice. PLoS One. 2015;10(2).

47. McGill BE, Bundle SF, Yaylaoglu MB, Carson JP, Thaller C, Zoghbi HY. Enhanced anxiety and stress-induced corticosterone release are associated with increased Crh expression in a mouse model of Rett syndrome. Proc Natl Acad Sci U S A. 2006;103(48).

48. Mcclennen SJ, Cortright DN, Seasholtz AF. Regulation of pituitary corticotropin-releasing hormone-binding protein messenger ribonucleic acid levels by restraint stress and adrenalectomy. Endocrinology. 1998;139(11):4435–41.

49. Hering H, Sheng M. Dentritic spines: structure, dynamics and regulation. Nat Rev Neurosci. 2001;

50. Segal M. Dendritic spines and long-term plasticity. Nat Rev Neurosci. 2005 Apr;6(4):277–84.

51. Droste SK, De Groote L, Atkinson HC, Lightman SL, Reul JMHM, Linthorst ACE. Corticosterone levels in the brain show a distinct ultradian rhythm but a delayed response to forced swim stress. Endocrinology. 2008;149(7):3244–53.

52. Qian X, Droste SK, Gutièrrez-Mecinas M, Collins A, Kersanté F, Reul JMHM, et al. A rapid release of corticosteroid-binding globulin from the liver restrains the glucocorticoid hormone response to acute stress. Endocrinology. 2011;152(10):3738–48.

53. Salvatore M, Wiersielis KR, Luz S, Waxler DE, Bhatnagar S, Bangasser DA. Sex differences in circuits activated by corticotropin releasing factor in rats. Horm Behav. 2018;

54. Boutillier AL, Sassone-Corsi P, Loeffler JP. The Protooncogene c-fos is induced by corticotropin-releasing factor and stimulates proopiomelanocortin gene transcription in pituitary cells. Mol Endocrinol. 1991;

55. Dedic N, Kühne C, Gomes KS, Hartmann J, Ressler KJ, Schmidt M V., et al. Deletion of CRH From GABAergic Forebrain Neurons Promotes Stress Resilience and Dampens Stress-Induced Changes in Neuronal Activity. Front Neurosci. 2019;

56. Henckens MJAG, Printz Y, Shamgar U, Dine J, Lebow M, Drori Y, et al. CRF receptor type 2 neurons in the posterior bed nucleus of the stria terminalis critically contribute to stress recovery. Mol Psychiatry. 2017 Dec 23;22(12):1691–700.

57. Miyata M, Okada D, Hashimoto K, Kano M, Ito M. Corticotropin-releasing factor plays a permissive role in cerebellar long-term depression. Neuron. 1999;22(4):763–75.

58. Tak PW, Howland JG, Robillard JM, Ge Y, Yu W, Titterness AK, et al. Hippocampal long-term depression mediates acute stress-induced spatial memory retrieval impairment. Proc Natl Acad Sci U S A. 2007;104(27):11471–6.

59. Sumi T, Harada K. Mechanism underlying hippocampal long-term potentiation and depression based on competition between endocytosis and exocytosis of AMPA receptors. Sci Rep. 2020;

60. Lemon N, Manahan-Vaughan D. Dopamine D1/D5 receptors contribute to de novo hippocampal LTD mediated by novel spatial exploration or locus coeruleus activity. Cereb Cortex. 2012;22(9):2131–8.

61. Gray EG. Problems of Interpreting the Fine Structure of Vertebrate and Invertebrate Synapses. In 1966. p. 139–70.

62. Bloom FE, Aghajanian GK. Fine structural and cytochemical analysis of the staining of synaptic junctions with phosphotungstic acid. J Ultrasructure Res. 1968;

63. Yuen EY, Liu W, Karatsoreos IN, Feng J, McEwen BS, Yan Z. Acute stress enhances glutamatergic transmission in prefrontal cortex and facilitates working memory. Proc Natl Acad Sci U S A. 2009;

64. Parihar VK, Hattiangady B, Kuruba R, Shuai B, Shetty AK. Predictable chronic mild stress improves mood, hippocampal neurogenesis and memory. Mol Psychiatry. 2011;

65. Adzic M, Djordjevic J, Djordjevic A, Niciforovic A, Demonacos C, Radojcic M, et al. Acute or chronic stress induce cell compartment-specific phosphorylation of glucocorticoid receptor and alter its transcriptional activity in Wistar rat brain. J Endocrinol. 2009;202(1):87–97.

66. Chen Y, Brunson KL, Adelmann G, Bender RA, Frotscher M, Baram TZ. Hippocampal corticotropin releasing hormone: Pre- and postsynaptic location and release by stress. Neuroscience. 2004;126(3):533–40.

67. Aguilar-Valles A, Sánchez E, De Gortari P, Balderas I, Ramírez-Amaya V, Bermúdez-Rattoni F, et al. Analysis of the stress response in rats trained in the water-maze: Differential expression of corticotropin-releasing hormone, CRH-R1, glucocorticoid receptors and brain-derived neurotrophic factor in limbic regions. Neuroendocrinology. 2006;82(5–6):306–19.

68. Jiang Z, Rajamanickam S, Justice NJ. CRF signaling between neurons in the paraventricular nucleus of the hypothalamus (PVN) coordinates stress responses. Neurobiol Stress. 2019 Nov;11:100192.

69. Doyon C, Samson P, Lalonde J, Richard D. Effects of the CRF1 receptor antagonist SSR125543 on energy balance and food deprivation-induced neuronal activation in obese Zucker rats. J Endocrinol. 2007;

70. Skórzewska A, Bidziński A, Hamed A, Lehner M, Turzyńska D, Sobolewska A, et al. The influence of CRF and α-helical CRF(9-41) on rat fear responses, c-Fos and CRF expression, and concentration of amino acids in brain structures. Horm Behav. 2008;

71. Gunn BG, Cox CD, Chen Y, Frotscher M, Gall CM, Baram TZ, et al. The endogenous stress hormone CRH modulates excitatory transmission and network physiology in hippocampus. Cereb Cortex. 2017;

72. Bajo M, Cruz MT, Siggins GR, Messing R, Roberto M. Protein kinase C epsilon mediation of CRF- and ethanol-induced GABA release in central amygdala. Proc Natl Acad Sci U S A. 2008;105(24):8410–5.

73. Hollrigel GS, Chen K, Baram TZ, Soltesz I. The pro-convulsant actions of corticotropin-releasing hormone in the hippocampus of infant rats. Neuroscience. 1998;84(1):71–9.

74. Giesbrecht CJ, Mackay JP, Silveira HB, Urban JH, Colmers WF. Countervailing modulation of Ihby neuropeptide Y and corticotrophin-releasing factor in basolateral amygdala as a possible mechanism for their effects on stress-related behaviors. J Neurosci. 2010;

75. Van Pett K, Viau V, Bittencourt JC, Chan RKW, Li HY, Arias C, et al. Distribution of mRNAs encoding CRF receptors in brain and pituitary of rat and mouse. J Comp Neurol. 2000;

76. Hiroi N, Wong ML, Licinio J, Park C, Young M, Gold PW, et al. Expression of corticotropin releasing hormone receptors type I and type II mRNA in suicide victims and controls. Mol Psychiatry. 2001;

77. Gunn BG, Baram TZ. Stress and Seizures: Space, Time and Hippocampal Circuits. Trends Neurosci. 2017 Nov;40(11):667–79.

78. Carboni L, Romoli B, Bate ST, Romualdi P, Zoli M. Increased expression of CRF and CRF-receptors in dorsal striatum, hippocampus, and prefrontal cortex after the development of nicotine sensitization in rats. Drug Alcohol Depend. 2018;

79. Smith GW, Aubry J-MM, Dellu F, Contarino A, Bilezikjian LM, Gold LH, et al. Corticotropin Releasing Factor Receptor 1–Deficient Mice Display Decreased Anxiety, Impaired Stress Response, and Aberrant Neuroendocrine Development. Neuron. 1998 Jun;20(6):1093–102.

80. Dedic N, Chen A, Deussing JM. The CRF Family of Neuropeptides and their Receptors - Mediators of the Central Stress Response. Curr Mol Pharmacol. 2018;11(1).

81. Bagosi Z, Balangó B, Pintér D, Csabafi K, Jászberényi M, Szabó G, et al. The effects of CRF and urocortins on the hippocampal glutamate release. Neurochem Int. 2015;

82. Bale TL, Contarino A, Smith GW, Chan R, Gold LH, Sawchenko PE, et al. Mice deficient for corticotropin-releasing hormone receptor-2 display anxiety-like behaviour and are hypersensitive to stress. Nat Genet. 2000;24(4):410–4.

83. Coste SC, Kesterson RA, Heldwein KA, Stevens SL, Heard AD, Hollis JH, et al. Abnormal adaptations to stress and impaired cardiovascular function in mice lacking corticotropin-releasing hormone receptor-2. Nat Genet. 2000;

